# Ras guanine nucleotide exchange factor RasGRP1 promotes acute inflammatory response and restricts inflammation-contributed cancer cell growth

**DOI:** 10.1101/2021.03.11.434895

**Authors:** Cong Wang, Xue Li, Changping Yu, Luoling Wang, Rilin Deng, Hui Liu, Zihao Chen, Yingdan Zhang, Suping Fan, Hungyu Sun, Haizhen Zhu, Jianli Wang, Songqing Tang

## Abstract

Acute inflammatory response needs to be tightly regulated for promoting the elimination of pathogens and preveting the risk of tumorigenesis, but the mechanism has not been fully elucidated. Here, we report that Ras guanine nucleotide releasing protein 1 (RasGRP1) plays a bifunctional regulator that promotes acute inflammation and inhibits inflammation-associated cancer. At the mRNA level, *RasGRP1* strengthens the inflammatory response by functioning as a competing endogenous RNA to specifically promote IL-6 expression by sponging let-7a. *In vivo* overexpression of the *RasGRP1* 3’ untranslated region significantly aggravated lipopolysaccharide-induced systemic inflammation and dextran sulphate sodium-induced colitis in *IL-6*^+/+^ mice but not in *IL-6*^-/-^ mice. At the protein level, RasGRP1 restricts the growth of inflammation-contributed cancer cells by impairing EGFR-SOS1-Ras-AKT signalling. Tumour patients with high RasGRP1 expression showed a better clinical outcome than those with low expression. Considering acute inflammation rarely leads to tumorigenesis, this work reveals that RasGRP1 is an essential bifunctional regulator for acute inflammatory response.

## Introduction

IL-6 is a key inflammatory cytokine in infection and cancer and was recently identified as a prominent target for clinical intervention ^1–4^. Almost all stromal cells and immune cells produce IL-6. Cytokines, bacteria and viruses are major inducers of IL-6 production ^5, 6^. IL-6 expression is relatively low in resting innate immune cells but is rapidly elevated in activated innate immune cells triggered by microbial components such as lipopolysaccharide (LPS) ^7^. In maintainance of IL-6 level under normal physiological conditions, microRNA (miRNA) let-7 which is abundantly expressed in resting innate immune cells represses the expression of IL-6 by binding its 3’ untranslated region (UTR) ^8–10^. However, let-7 expression shows only a slight downregulation in activated innate immune cells ^10^. Thus, the rapid-responsed inhibitory effect of let-7 on IL-6 of activated innate immune cells, especially during the stage of acute inflammatory response, may be protential druggble targets for inflammatory disease.

miRNAs are a class of small noncoding RNAs that modulate gene expression though mRNA degradation or translational inhibition ^11^. As miRNA sponges, RNA transcripts show crosstalk and regulate each other by competing for their shared miRNAs, known as competing endogenous RNAs (ceRNAs) ^12, 13^. Based on target competition, ceRNAs exhibit positively correlated and temporal, spatial and disease-specific expression patterns ^13^. Recent reports have proven that mRNAs, long noncoding RNAs, pseudogenes and circular RNAs can act as ceRNAs to affect the progression of cancer or neurological diseases ^14–19^. Interestingly, miRNA targeted competition by sponges occurs in a threshold-like manner ^20^. Recently, it had been demonstrated that ceRNA mediated derepression miRNA targets was observed just after exceeding a high threshold of added ceRNA ^21^. Coincidentally, the expression of inflammatory cytokines, such as IL-6, is substantially upregulated in acute inflammatory response. So, the cross-talk between ceRNAs and the expression of inflammatory cytokines at acute inflammatory response remains to be further investigated.

Ras guanine nucleotide-releasing protein 1 (RasGRP1), a member of the RasGRP family, is a guanine nucleotide exchange factor for Ras ^22, 23^. The catalytic region of RasGRP1 consists of a Ras exchange motif (REM) and a cell division cycle 25 (CDC25) domain, and the regulatory region of RasGRP1 contains two EF hands (calcium binding) and a C1 domain (DAG binding) ^22, 24^. RasGRP1 is highly expressed in T cells and can also be detected in B cells, mast cells, natural killer cells, neuronal cells and activated macrophages ^25–30^. Previous studies had found that RasGRP1 is critical in mediating Ras/Erk activation, thus promotes the positive and negative selection of T cells ^31–33^. Recently, increasing evidence has shown that RasGRP1 plays an important role in many human diseases, especially inflammatory diseases such as systemic lupus erythematosus, type 2 diabetes, and cancer ^34–38^. Although the function of RasGRP1 has been extensively investigated in T cells, its role in innate immune cells, especially in macrophages, is still unclear. Unveiling of the role of RasGRP1 in innate immunity may provide new insight into the pathophysiology and underlying mechanism of inflammatory diseases.

In the current study, we investigated the role of RasGRP1 in innate immunity and inflammatory diseases. We found that RasGRP1 specifically derepresses the expression of IL-6 by competing with let-7a during the acute inflammatory response. Our data elucidated a previously uncharacterized coding-independent role of RasGRP1 in innate immunity, providing insight into ceRNA modulation of acute inflammatory response, and showed that RasGRP1 is an important regulator for acute inflammation to promote the production of the proinflammtory cytokine IL-6 and decrease the probability of inflammation-associated cancer in a protein-independent or protein-dependent manner, respectively, which may explain why acute inflammation rarely leads to tumorigenesis.

## Results

### RasGRP1 and IL-6 are both regulated by let-7a

Previous studies have revealed that let-7 can directly inhibit the expression of IL-6 by binding its 3’UTR ^8, 9^. We examined the expression profiles of let-7a, let-7b, let-7c, let-7d, let-7e, let-7f, let-7g, let-7i and let-7k in mouse peritoneal macrophages and found that let-7a showed the highest expression level in these cells (Fig. 1a), suggesting that let-7a may be the major let-7 family member that regulates the expression of IL-6 in peritoneal macrophages. To confirm that let-7a inhibits the expression of IL-6, we cotransfected IL-6 expression plasmids with synthetic let-7a mimics and/or let-7a inhibitor into HEK293 cells. The result showed that let-7a mimics inhibited the expression of IL-6 by binding the *IL-6* 3’UTR, which was reversed by let-7a inhibitor (Fig. 1b). To determine whether let-7a is a critical mediator inhibiting IL-6 expression in macrophages, we overexpressed or silenced let-7a in macrophages by transfecting cells with synthetic let-7a mimics or let-7a inhibitor, respectively. Quantitative RT-PCR analysis showed that the *IL-6* mRNA levels were barely unchanged in the LPS-stimulated macrophages transfected with let-7a mimics or inhibitor (Fig. 1c). The IL-6 protein levels decreased in the let-7a mimic-transfected cells and increased in the let-7a inhibitor-transfected cells compared with the control cells after stimulation with LPS (Fig. 1d). These results suggested that IL-6 expression could be inhibited by let-7a via translational inhibition but not via mRNA degradation in macrophages.

**Fig. 1.**
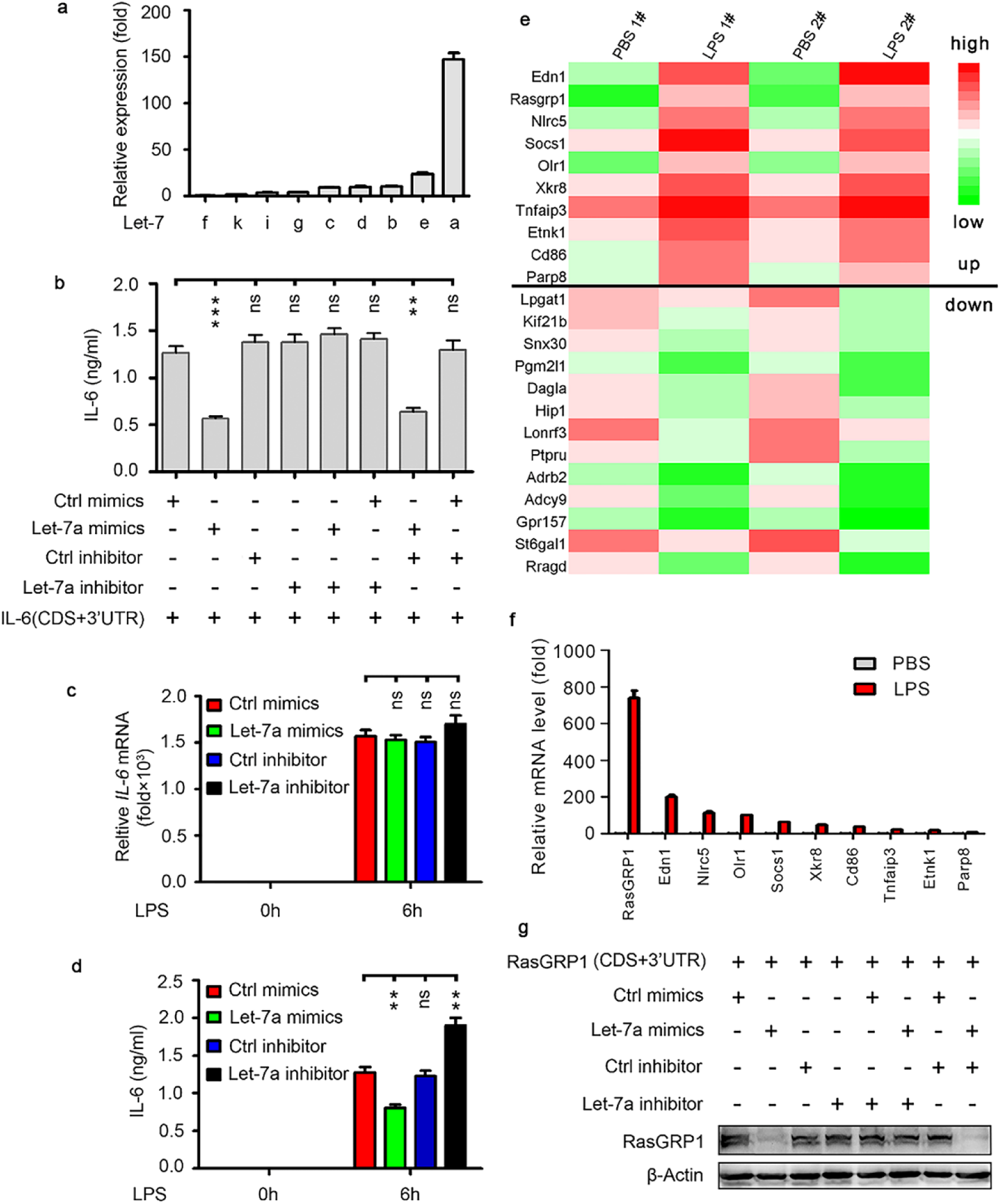
RasGRP1 and IL-6 are both regulated by let-7a. **a** Q-PCR analysis of microRNA Let-7 isoforms expression in peritoneal macrophages. **b** ELISA quantification of IL-6 in supernatants of HEK293 cells transfected with indicated molecules for 24h. **c** Q-PCR analysis of IL-6 mRNA expression in peritoneal macrophages transfected with Let-7a mimics, Let-7a inhibitor or matched control and 24h later treated with 100 ng/ml LPS for 6h. **d** ELISA quantification of IL-6 in supernatants of peritoneal macrophages treated as in (**c**). **e** Microarray analysis of mouse peritoneal macrophages treated with 100 ng/ml LPS. Shown is the heat map of the predicted target-genes of microRNA Let-7 upregulated or downregulated in macrophages following LPS treatment for 6h. Full list of diferentially expressed genes available at GEO (accession GSE165800). **f** Q-PCR analysis of predicted target-genes of microRNA Let-7 in mouse peritoneal macrophages treated with 100 ng/ml LPS. **g** Immunoblot analysis of RasGRP1 and β-actin in lysates of HEK293 cells transfected with indicated molecules for 24h. Data are shown as mean ± S.E.M. (**a-d, f, g**) of one representative experiment. Similar results were obtained in three independent experiments. ns, not significant; ** *p*<0.01.

To identify the possible targets of let-7a that might compete with IL-6 in the innate immune response, gene expression profiles in mouse peritoneal macrophages stimulated with LPS was further analysed. Focusing on the expression of 1046 putative let-7 target genes predicted by the TargetScan algorithm (http://www.targetscan.org), 23 putative let-7 target genes, including 10 upregulated genes, showed more than 5-fold changes in the LPS-stimulated macrophages (Fig. 1e). To verify the results of the gene expression profiles, 10 of upregulated genes were further quantified by qPCR and results showed that *RasGRP1* was the top upregulated gene among these 10 genes (Fig. 1f), indicating that *RasGRP1* might effectively sponge let-7a. To confirm that RasGRP1 was regulated by let-7a, we cotransfected the expression plasmids of *RasGRP1* with synthetic let-7a mimics and/or let-7a inhibitor into HEK293 cells and found that the let-7a mimics inhibited the expression of RasGRP1 by binding its 3’UTR and that the let-7a inhibitor reversed the effects of the let-7a mimics (Fig. 1g). In summary, these results indicated that IL-6 and RasGRP1 are both regulated by let-7a.

### *RasGRP1* is coexpressed with *IL-6* and *RasGRP1* silencing selectively inhibits IL-6 protein levels during acute inflammatory response

To determine whether *RasGRP1* is coexpressed with *IL-6* during acute inflammatory response, we treated mouse peritoneal macrophages with gradient doses of LPS and observed that *RasGRP1* and *IL-6* were significantly upregulated in a dose-dependent manner (Fig. 2a). A similar phenomenon was observed when peritoneal macrophages were stimulated by gradient doses of poly(I:C) (Fig. 2b) or CpG ODN (Fig. 2c). In addition, we stimulated mouse bone marrow-derived macrophages with gradient doses of LPS, poly (I:C) or CpG ODN. We also found that *RasGRP1* and *IL-6* were significantly upregulated in a dose-dependent manner (Fig. S1a-c). These results suggest that *RasGRP1* may be coexpressed with *IL-6*.

**Fig. 2.**
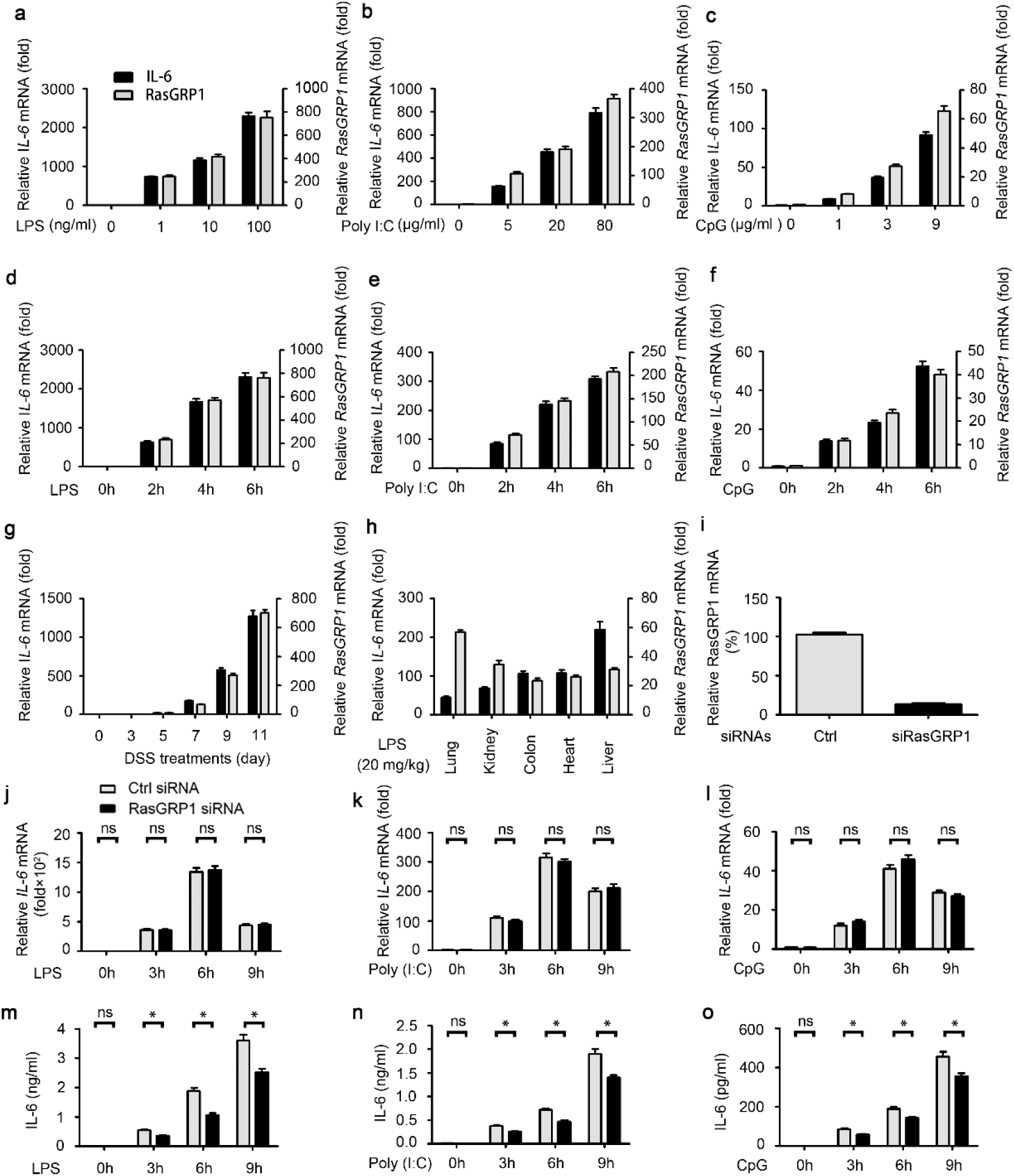
RasGRP1 co-expresses with IL-6 and RasGRP1 silence selectively inhibits IL-6 protein levels in acute inflammatory response. **a-c** Quantitative PCR (Q-PCR) analysis of IL-6 and RasGRP1 mRNA expression in peritoneal macrophages treated with different doses of LPS (0, 1, 10 or 100 ng/ml) (**a**), Poly (I:C) (0, 5, 20 or 80 μg/ml) (**b**), or CpG ODN (0, 1, 3 or 9 μg/ml) (**c**) for 6h. **d-f** Q-PCR analysis of IL-6 and RasGRP1 mRNA expression in peritoneal macrophages treated with 100 ng/ml LPS (**d**), 10 μg/ml Poly (I:C) (**e**), or 5 μg/ml CpG ODN (**f**) for the indicated hours. **g** Q-PCR analysis of IL-6 and RasGRP1 mRNA expression in intestinal epithelial cells isolated from DSS treated mice on the indicated days. **h** Q-PCR analysis of IL-6 and RasGRP1 mRNA expression in organs from mice injected intraperitoneally with LPS (15 mg per kg body weight) for 6h. **i** Q-PCR analysis of RasGRP1 mRNA expression in peritoneal macrophages 48h after transfection with RasGRP1 siRNA. **j-l** Q-PCR analysis of IL-6 mRNA expression in peritoneal macrophages transfected as in (**a**) and 48h later treated with 100 ng/ml LPS (**j**), 10mg/ml Poly (I:C) (**k**) or 5mM of CpG ODN (**l**) for the indicated hours. **m-o** ELISA quantification of IL-6 in supernatants of macrophages treated as in (**j-l**). Data are representative of three independent experiments with similar results (means ± S.D. in **a-g** and **i-n**). ns, not significant; * *p*<0.05.

To confirm the coexpression of *RasGRP1* and *IL-6*, we treated mouse peritoneal macrophages with 100 ng/ml LPS for different durations and found that *RasGRP1* and *IL-6* were quickly and significantly upregulated in a time-dependent manner (Fig. 2d). Similar results were obtained when peritoneal macrophages were stimulated with 10 μg/ml poly (I:C) (Fig. 2e) or 5 μg/ml CpG ODN (Fig. 2f) for different durations. In addition, we stimulated mouse bone marrow-derived macrophages with 100 ng/ml LPS, 10 μg/ml poly (I:C) or 5 μg/ml CpG ODN for different durations. We also observed that *RasGRP1* and *IL-6* were quickly and significantly upregulated in a time-dependent manner (Fig. S1d-f).

Next, we examined the coexpression of *RasGRP1* and *IL-6* in intestinal epithelial cells isolated from the inflamed colons of C57B/6 mice. We found that *RasGRP1* and *IL-6* were significantly upregulated in these cells in the days after the mice were fed 2.5% DSS (Fig. 2g). We also examined the coexpression of *RasGRP1* and *IL-6* in the cells isolated from the lung, kidney, colon, heart or liver of the C57B/6 mice challenged with LPS. We observed that both *RasGRP1* and *IL-6* were significantly upregulated in LPS-induced systemic inflammation (Fig. 2h). Together, these data indicated that *RasGRP1* is coexpressed with *IL-6* in the acute inflammation and inflammation-associated diseases.

To ascertain the role of RasGRP1 in acute inflammatory response, we silenced RasGRP1 in mouse peritoneal macrophages (Fig. 2i) and found that siRNA knockdown of RasGRP1 did not affect the mRNA expression of cytokines and chemokines in peritoneal macrophages stimulated with LPS, poly (I:C) or CpG ODN (Fig. 2j-l and Fig. S2a-o). Consistent with the mRNA expression of cytokines and chemokines, the activation of ERK, JNK and P38 MAPKs, as well as the IKKα/β-IκBα pathway, was not affected by RasGRP1 knockdown after LPS treatment (Fig. S2p). Interestingly, IL-6 secretion was significantly decreased when we stimulated RasGRP1-silenced peritoneal macrophages with LPS, poly (I:C) or CpG ODN (Fig. 2m-o). In contrast, no difference in TNF-α secretion was observed between the RasGRP1-silenced macrophages and the control macrophages (Fig. S2q-s). Taken together, these results showed that RasGRP1 selectively promote IL-6 protein levels in acute inflammatory response.

In human peripheral blood MDMs, we also examined the coexpression of *RasGRP1* and *IL-6*. We found that *RasGRP1* and *IL-6* were also significantly upregulated in the MDMs stimulated with gradient doses of LPS (Fig. S3a), poly (I:C) (Fig. S3b) or CpG ODN (Fig. S3c). Similar results were obtained when MDMs were stimulated with 100 ng/ml LPS (Fig. S3d), 10 μg/ml poly (I:C) (Fig. S3e) or 5 μg/ml CpG ODN (Fig. S3f) for different durations.

Next, we also silenced RasGRP1 in human MDMs (Fig. S3g) and found that siRNA knockdown of RasGRP1 did not affect the mRNA expression of cytokines and chemokines in the human MDMs stimulated with LPS, poly (I:C) or CpG ODN (Fig. S3h-m). Consistent with the results in mouse peritoneal macrophages, IL-6 secretion was significantly decreased when we silenced RasGRP1 in the human MDMs stimulated with LPS, poly (I:C) or CpG ODN (Fig. S3n). However, TNF-α secretion was not affected (Fig. S3o). Taken together, these results showed that *RasGRP1* also selectively promote IL-6 protein levels in human MDMs during acute inflammatory response.

### The *RasGRP1* 3’UTR enhances IL-6 protein levels by sponging let-7a

To verify whether *RasGRP1* compete with let-7a for *IL-6*, we cotransfected the *IL-6* expression vector and let-7a mimics with the *RasGRP1* CDS and/or 3’UTR expression vector into HEK293 cells and found that the *RasGRP1* 3’UTR, but not the *RasGRP1* CDS, promoted the expression of IL-6 by competing with let-7a for IL-6 (Fig. 3a). To determine which position of the *RasGRP1* 3’UTR bind to let-7a, three candidate let-7a-sponging RNAs with 30 bp in lenth was synthesised according to the nearby sequence of the potential let-7 binding site on the *RasGRP1* 3’UTR (Fig. 3b). Confocal microscopy showed that only the sponging RNA candidate containing the complementary sequence to the seed sequence(Fig. S4), which is important for the binding of the miRNA to mRNA, bind to let-7a (Fig. 3c). Next, the effect of the three let-7a-sponging RNA candidates was examined by quantitative RT-PCR and results showed that the let-7a-sponging RNAs had no significant effect on the *IL-6* mRNA levels in the LPS-stimulated macrophages regardless presense of the let-7a mimic (Fig. 3d). ELISA assay showed that only the sponging RNA candidate containing the complementary sequence to the seed sequence could promote the IL-6 protein levels by competing with IL-6 in macrophages after stimulation with LPS (Fig. 3e). These data suggested that the sponging RNA candidate containing the complementary sequence to the seed sequence, similar to the *RasGRP1* 3’UTR, positively regulate the production of IL-6 protein in macrophages during acute inflammatory response.

**Fig. 3.**
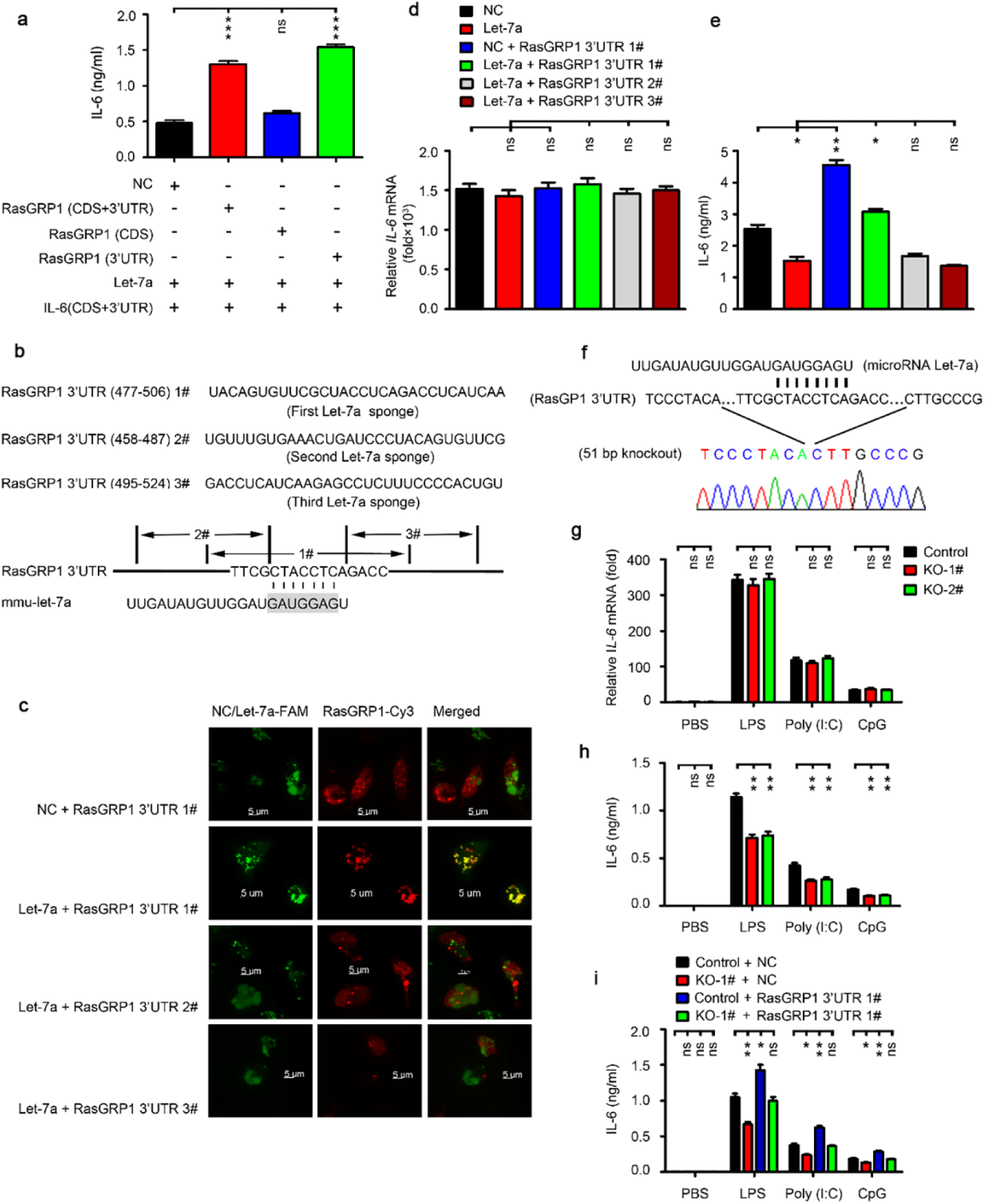
The 3’UTR of RasGRP1 enhances IL-6 protein levels by sponging let-7a. **a** ELISA quantification of IL-6 in supernatants of HEK293 cells <u>transfected</u> with plasmids expressing the CDS and/or 3’UTR of RasGRP1 and IL-6 and transfected with let-7a mimics for 24 h. **b** A schematic diagram showing the sequence of the sponge RNA designed according to the sequence of the RasGRP1 3’UTR. **c** Representative confocal microscopic image of peritoneal macrophages cotransfected with NC/let-7a-FAM (green) and its sponge RNA RasGRP1 3’UTR #1, #2 or #3-Cy3 (red) designed as in (**b**). **d** qPCR analysis of IL-6 mRNA expression in peritoneal macrophages transfected with Let-7a and/or RasGRP1 3’UTR #1, #2 or #3 and 24 h later treated with 100 ng/ml LPS for 6 h. **e** ELISA quantification of IL-6 in the supernatants of peritoneal macrophages treated as in (**d**). **f** DNA sequencing of PCR products confirmed the targeted mutation at the 3’UTR of the RasGRP1 gene in the RAW264.7 macrophage clone. The mutation contains a 51-bp deletion, which included the binding site of let-7a. **g** qPCR analysis of IL-6 mRNA expression in the RAW264.7 macrophage clones KO-1# or KO-2# with a 51-bp deletion as in (**f**) and treated with LPS, poly (I:C) or CpG for 6 h. **h** ELISA quantification of IL-6 in the supernatants of the RAW264.7 macrophages treated as in (**G**). **i** ELISA quantification of IL-6 in the supernatants of the RAW264.7 macrophage clone KO-1# transfected with RasGRP1 3’UTR #1 and 24 h later treated with LPS, poly (I:C) or CpG for 6 h. Data are shown as the mean ± S.E.M. Similar results were obtained in three independent experiments. ns, not significant; \**p*<0.05.; \*\**p*<0.01.

To provide further evidence, two sgRNAs and three genotyping primers were designed to deplete the let-7a binding site on the 3’UTR of *RasGRP1* in RAW264.7 cells via the genome-editing technique (Fig. S5a). Five candidate *RasGRP1* 3’UTR mutant clones in RAW264.7 cells were confirmed by PCR screening (Fig. S5b). DNA sequencing futher confirmed two positive clones among the five candidate mutant clones (Fig. 3f). In the two clones with mutations, we observed that the *IL-6* mRNA levels were not affected, but the IL-6 protein levels were significantly decreased after LPS, poly (I:C) or CpG ODN treatment (Fig. 3g and h). To exclude off-target effects, we transfected the sponging RNA candidate containing the complementary sequence to the seed sequence into the positive clones. We found that the sponging RNA candidate containing the complementary sequence to the seed sequence rescued the effect of the *RasGRP1* 3’UTR mutation on the IL-6 protein levels after TLR3/4/9 agonist treatments (Fig. 3i). Together, these results suggest that the *RasGRP1* 3’UTR sponge let-7a to promote IL-6 protein levels during acute inflammatory response.

### *RasGRP1* mRNA 3’UTR aggravates LPS-induced systemic inflammation in *IL-6*^+/+^ mice

To investigate the role of the *RasGRP1* 3’UTR *in vivo*, we synthesized cholesterol-conjugated let-7a-sponging RNA and let-7a mimics. By intraperitoneal injection of cholesterol-conjugated let-7a-sponging RNA and let-7a mimics, we found that the sponging RNA candidate containing the complementary sequence to the seed sequence also effectively bind to let-7a mimics *in vivo* (Fig. 4a). In response to LPS, the mice intravenously injected with the cholesterol-conjugated let-7a mimics produced less IL-6 than did the control mice, and the sponging RNA candidate containing the complementary sequence to the seed sequence reversed these effects of the let-7a mimics (Fig. 4b). After lethal challenge with LPS, the *IL-6*^+/+^ mice, but not the *IL-6*^-/-^ mice, intravenously injected with cholesterol-conjugated Let-7a displayed substantially prolonged survival, whereas the *IL-6*^+/+^ mice treated with let-7a-sponging RNA displayed strongly reduced survival (Fig. 4c and d). Furthermore, after intraperitoneal injection of the cholesterol-conjugated let-7a mimics, we observed milder inflammatory infiltration in the mouse lungs, but intraperitoneal injection of the cholesterol-conjugated let-7a-sponging RNA caused more severe inflammatory infiltration in the mouse lungs, and the sponging RNA candidate containing the complementary sequence to the seed sequence reversed the effects of the let-7a mimics (Fig. 4e). IHC staining showed that the let-7a mimics inhibited the production of IL-6 (Fig. 4f) in the mouse lungs, but the sponging RNA candidate containing the complementary sequence to the seed sequence promoted productions. All these results suggested that the 3’UTR of *RasGRP1* promotes LPS-induced systemic inflammation by competing for let-7a with IL-6 and enhancing the production of IL-6.

**Fig. 4.**
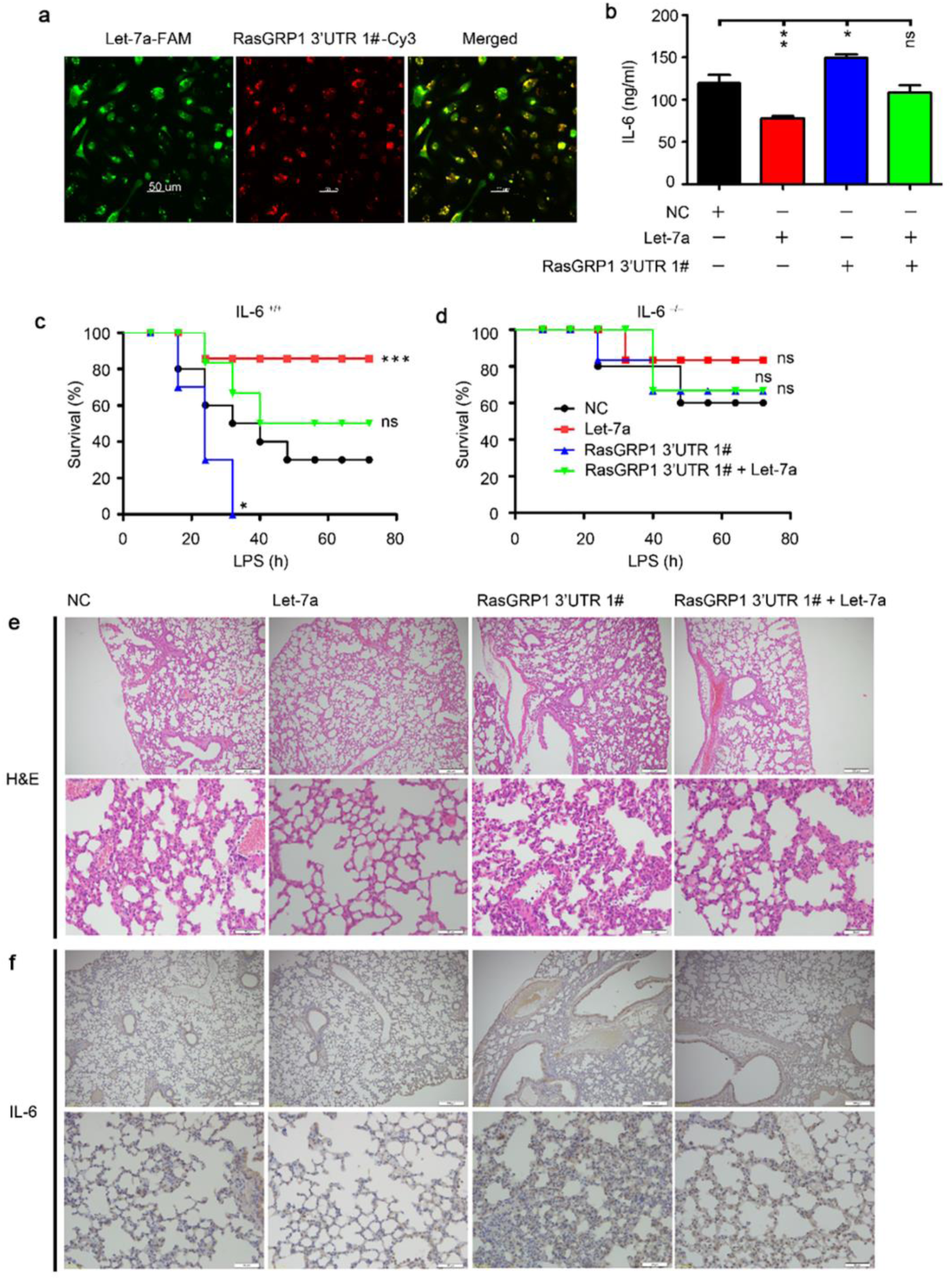
The 3’UTR of RasGRP1 aggravates LPS-induced systemic inflammation in IL-6^+/+^ mice. **a** Representive confocal microscopic image of peritoneal macrophages isolated from the mice treated by intraperitoneal injection of cholesterol-conjugated microRNA Let7a-FAM (Green) and its sponge RNA RasGRP1 3’UTR 1#-Cy3 (Red). **b** ELISA quantification of IL-6 in serum from IL-6^+/+^ mice treated by intravenous injection of cholesterol-conjugated microRNA Let-7a and/or its sponge RNA RasGRP1 3’UTR 1#, and 24 h later injected intraperitoneally with LPS for 2 h. **c, d** Survival of IL-6^+/+^ mice (**c**) and IL-6^−/−^ mice (**d**) treated as in (**b**) and monitored every hour after lethal challenge with LPS. **e** Hematoxylin and eosin staining of lungs from IL-6^+/+^ mice (n=6 per group) treated by intravenous injection of cholesterol-conjugated microRNA Let-7a and/or its sponge RNA RasGRP1 3’UTR 1#, and 24 h later injected intraperitoneally with LPS for 8 h. Scale bars represent 200 μm (top) and 50 μm (bottom). **f** Immunohistochemistry staining for IL-6 protein in the lungs of IL-6^+/+^ mice (n=6 per group) treated as in (**e**). Scale bars represent 200 μm (top) and 50 μm (bottom). Data are shown as mean ± S.E.M. All experiments were repeated three times. ns, not significant; * *p*<0.05.; ** *p*<0.01.

### The *RasGRP1* 3’UTR aggravates DSS-induced colitis in *IL-6*^+/+^ mice

Our data indicated that the *RasGRP1* 3’UTR promotes IL-6 protein production in TLR-induced inflammatory responses both *in vitro* and *in vivo*. To further investigate the physiological significance of these observations, we analysed the role of the *RasGRP1* 3’UTR in the development of DSS-induced colitis. After DSS treatment, we found that body weight loss was significantly retarded in the *IL-6*^+/+^ mice but not in the *IL-6*^-/-^ mice intravenously injected with cholesterol-conjugated let-7a (Fig. 5a and b). However, body weight loss was significantly faster in the *IL-6*^+/+^ mice intravenously injected with cholesterol-conjugated let-7a-sponging RNA (Fig. 5a). Moreover, the mice intravenously injected with cholesterol-conjugated let-7a mimic showed less severe rectal bleeding and colon shortening (Fig. 5c and d), lower serum IL-6 levels (Fig. 5e and f), less destruction of the bowel wall and less inflammatory cell infiltration than the control mice (Fig. 5g). The mice intravenously injected with cholesterol-conjugated let-7a-sponging RNA showed more severe rectal bleeding and colon shortening (Fig. 5c and d), higher serum IL-6 levels (Fig. 5e and f), more destruction of the bowel wall and more inflammatory cell infiltration than the control mice (Fig. 5g). When the mice were intravenously injected with cholesterol-conjugated let-7a mimic and let-7a-sponging RNA, the phenotype of these mice was similar to that of the control mice (Fig. 5). Collectively, our data suggest that the 3’UTR of *RasGRP1* aggravates the development of colitis by promoting the production of IL-6 protein.

**Fig. 5.**
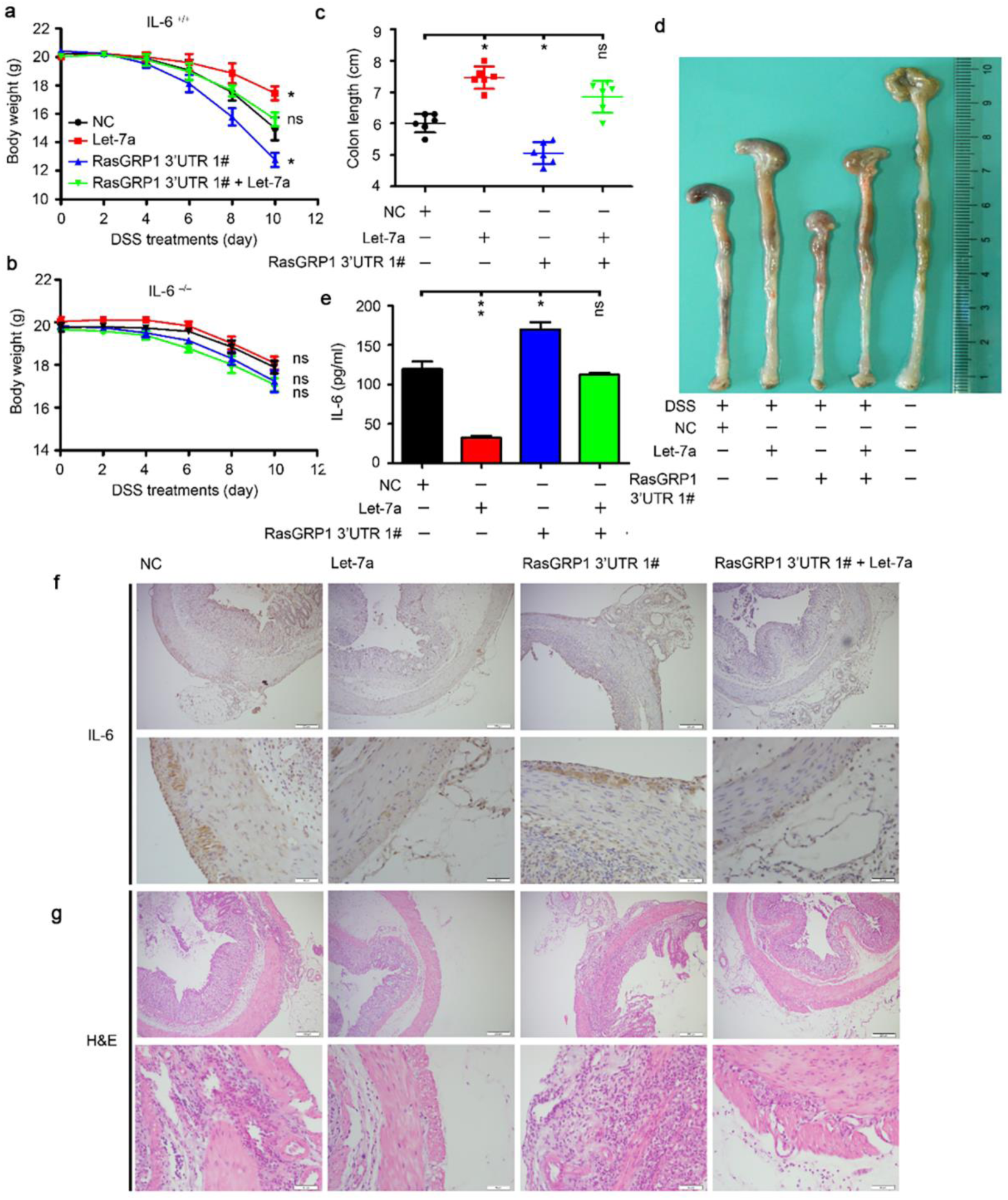
The 3’UTR of RasGRP1 aggravates dextrane sulphate sodium-induced colitis in IL-6^+/+^ mice. **a, b** Weight change over time of IL-6^+/+^ mice (**a**) and IL-6^−/−^ mice (**b**) (n=8 per group) treated by intraperitoneal injection of cholesterol-conjugated microRNA Let-7a and/or its sponge RNA RasGRP1 3’UTR 1#, and were fed with 2.5% DSS in water for 7 days. **c** ELISA quantification of IL-6 in serum from IL-6^+/+^ mice (n=6 per group) treated as in (**a**) on day 10 after DSS treatments. **d** Gross appearances of representative colons collected from IL-6^+/+^ mice (n=6 per group) treated as in (**a**) on day 10 after DSS treatments. **e** The length of colons from the IL-6^+/+^ mice (n=6 per group) treated as in (**a**) was measured on day 10 after DSS treatments. **f** Immunohistochemistry staining for IL-6 protein in the colons of IL-6^+/+^ mice (n=6 per group) treated as in (**a**) on day 10 after DSS treatments. Scale bars represent 200 μm (top) and 50 μm (bottom). **g** Hematoxylin and eosin staining of colons from the IL-6^+/+^ mice (n=6 per group) treated as in (**f**). Scale bars represent 200 μm (top) and 50 μm (bottom).Data are shown as mean ± S.E.M. All experiments were repeated three times. ns, not significant; \**p*<0.05.; \*\**p*<0.01.

### RasGRP1 restricts the growth of inflammation-contributed cancer cells

Previous study has revealed that IL-6 signaling contributes to the malignant progression of liver cancer progenitors^39^, and a recent study reported that RasGRP1 upregulation is associated with a better prognosis in colorectal cancer patients by limiting proliferative EGFR-SOS1-Ras-ERK signals ^40^. Therefore, we speculate that RasGRP1 may inhibit the development of liver cancer by limiting EGFR-SOS1-Ras-ERK signalling to prevent the tumour-promotion effect of IL-6 during acute inflammatory response. To confirm our hypothesis, we tested the expression levels of *RasGRP1*, *SOS1*, *SOS2*, *EGF* and *EGFR* in Huh7 and HepG2 cells and found that *EGF* and *EGFR* had higher expression in Huh7 cells than in HepG2 cells (Fig. 6a). Then, we analysed the role of RasGRP1 in Huh7 cells treated with 10 ng/ml EGF and found that RasGRP1 inhibited the proliferation of Huh7 cells (Fig. 6b) mainly by limiting EGFR-SOS1-Ras-AKT signal (Fig. 6c and d). Additionally, scratch wounding assays further showed that overexpression of RasGRP1 inhibited the proliferation of Huh7 cells (Fig. 6e and f). Next, we performed immunohistochemical staining for RasGRP1 on a human liver cancer tissue microarray. Representative images of liver cancer tissue samples with high, moderate or weak (according to the IHC score: weak⩽1; 1<moderate⩽2; 2=high⩽3) RasGRP1 expression are shown in (Fig. 6g). Interestingly, only 22.2% of the liver cancer tissue samples had high RasGRP1 expression compared to the percentages of moderate and weak RasGRP1 expression (40% and 37.8%, respectively) (Fig. 6h). However, 50% of the liver cancer tissue samples showed upregulation of RasGRP1 compared to the matched adjacent liver tissues (Fig. 6i). In the RasGRP1-upregulated tumour samples, the RasGRP1 IHC score was higher than that in RasGRP1-downregulated tumour samples (Fig. 6j). The liver cancer patients with upregulated RasGRP1 expression had a significantly smaller tumour size (Fig. 6k) and lower γ-glutamyl transferase (Fig. 6l) and longer total survival (Fig. 6m) than the liver cancer patients with downregulated RasGRP1 expression. Taken together, our results indicated that RasGRP1 protein can inhibit liver cancer by limiting EGFR-SOS1-Ras-AKT signal.

**Fig. 6.**
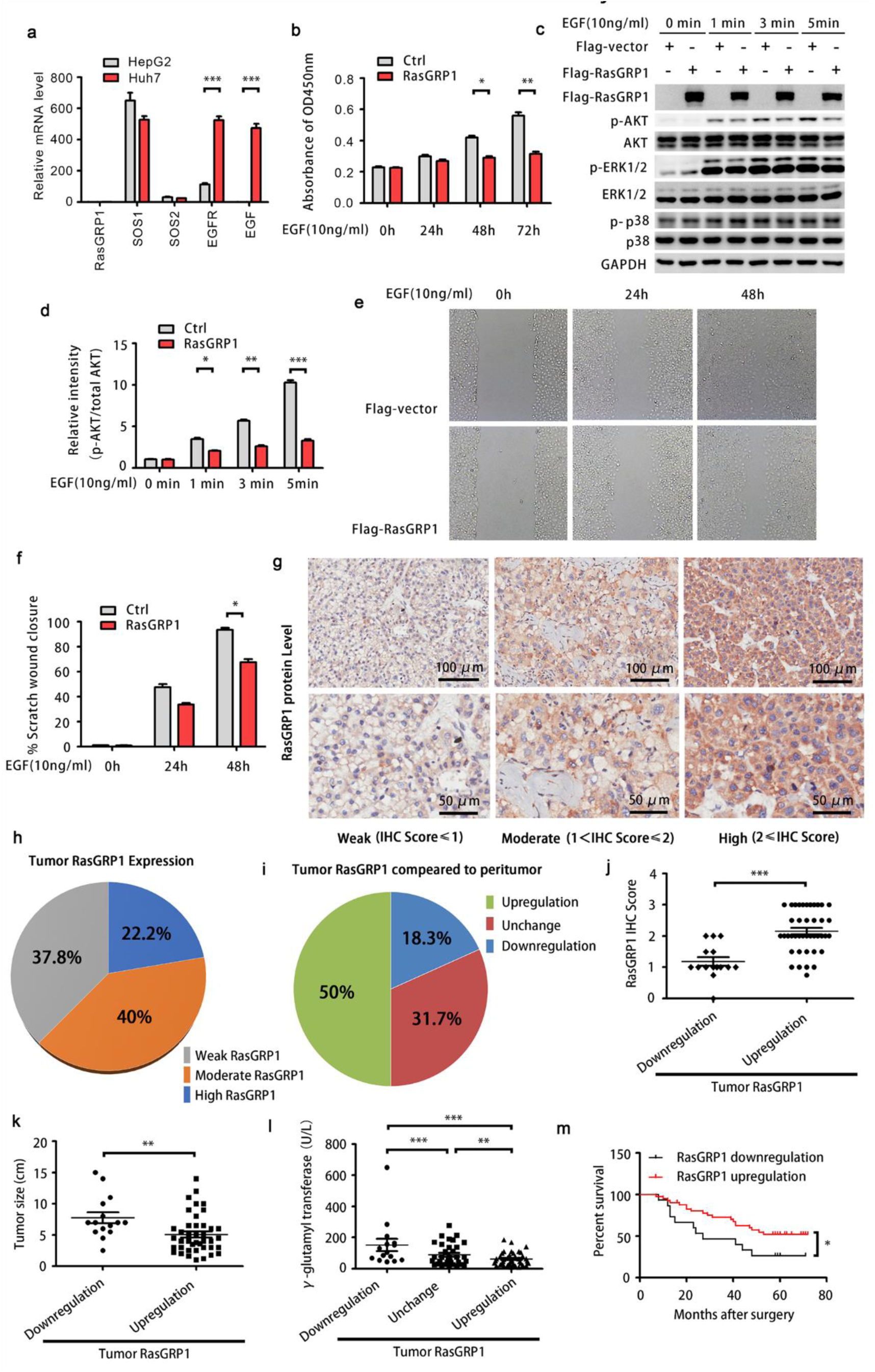
RasGRP1 affects the growth of liver cancer cells and the prognosis of liver cancer patients. **a** qPCR analysis of RasGRP1, SOS1, SOS2, EGF and EGFR in Huh7 and HepG2 cells. **b** CCK-8 assays were used to detect the role of RasGRP1 in the Huh7 cells treated with 10 ng/ml EFG. **c** Immunoblot analysis of the indicated molecules in lysates of the Huh7 cells treated with 10 ng/ml EFG for the indicated minutes. **d** The activation of AKT in (**c**) was quantified by determining the band intensity and calculated as phosphorylated AKT (Ser473) to total AKT. **e** Analysis of Huh7 cell migration by scratch wound closure assays. Images were acquired at the indicated minutes. **f** Quantitative analysis of (**e**). **g** Representative IHC staining showing high, moderate and weak RasGRP1 protein levels in a human liver cancer tissue microarray. **h** Pie chart of the IHC-determined RasGRP1 expression in tumour tissues according to (**g**). **i** Demographic graph of the liver cancer patients with RasGRP1 expression changes in tumour tissues compared to paired adjacent liver tissues. **j** The expression levels of RasGRP1 between liver cancer tissues (n = 41) and matched adjacent liver tissues (n = 15) were compared. **k**, **l** The association between RasGRP1 and tumour size (**k**) or γ-glutamyl transferase (**l**) was analysed. **m** Kaplan-Meier analysis of RasGRP1 expression and overall survival of the liver cancer patients. Data are representative of three independent experiments with similar results. * *p* <0.05; ** *p* <0.01; *** *p* <0.001.

To ascertain whether RasGRP1 protein restricts other cancers, We analysed the effect of RasGRP1 gene expression on the survival of multiple kinds of cancer patients according to the Kaplan-Meier Plotter database (http://kmplot.com/analysis/) and the existed data of the database shows that high expression of RasGRP1 positively correlated (Fig. 7) or no significantly correlated (Fig. S6a-k) with a improved overall survival of many kinds of cancer patients, and only in patients with kidney renal palillary cell carcinoma, high expression of RasGRP1 positively correlated with a poorer overall survival of patients (Fig. S6l). These results suggested that RasGRP1 protein might restrict the growth of multiple kinds of cancer cells.

**Fig. 7.**
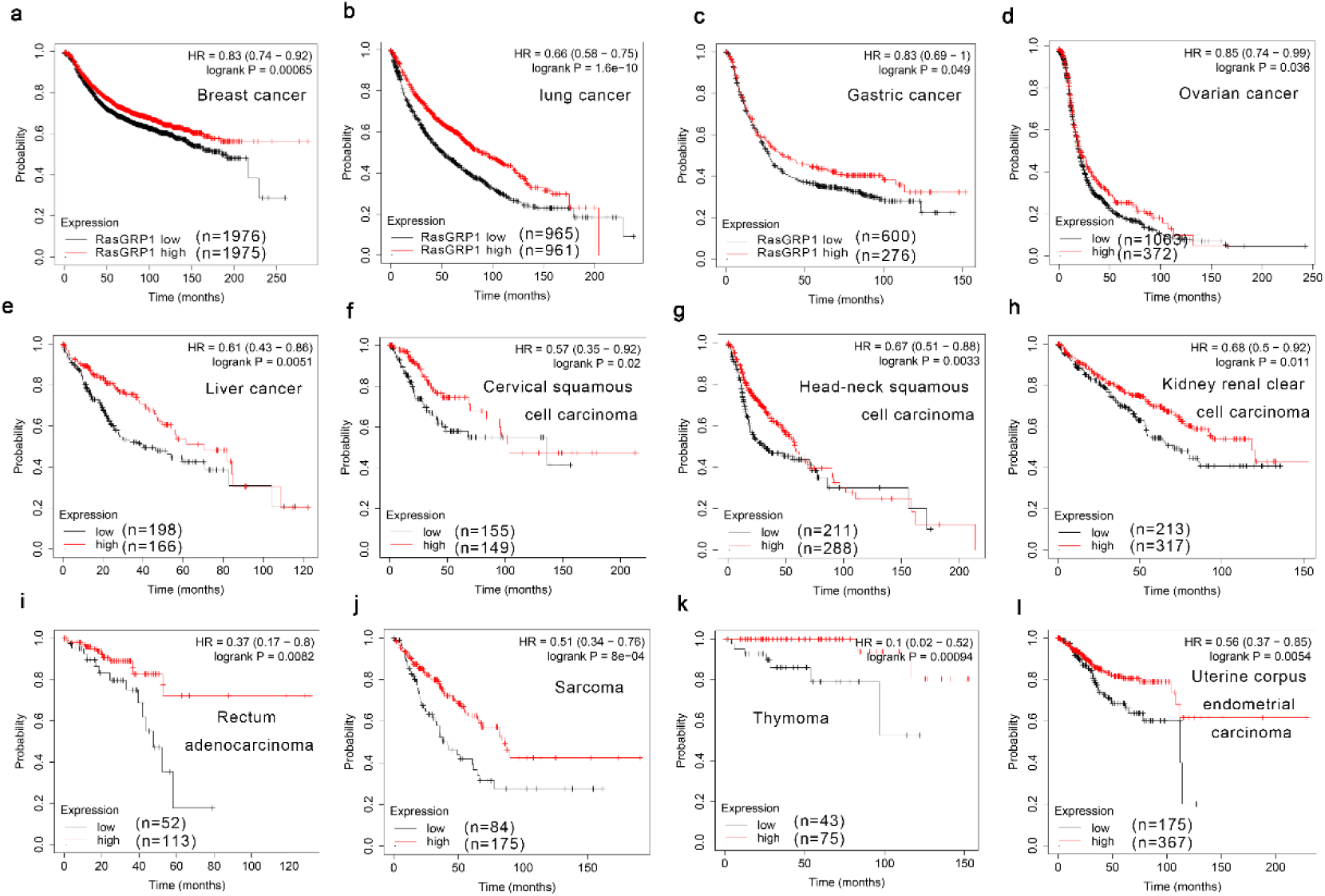
RasGRP1 expression correlates with the overall survival of cancer patients. **a-l** The survival curves from Kaplan-Meier plot profiles (http://kmplot.com/analysis/) for cancer patients stratified by high and low expression of RasGRP1.

## Discussion

In this study, we revealed that RasGRP1 has an important function in regulating acute inflammatory response. Acute inflammatory response is crucial for the efficient clearance of pathogens, but needs to be strictly regulated for preventing deleterious sideffects, such as development of tumours and autoimmune diseases ^9, 41^. Our study demonstrated that RasGRP1 is an key regulator in promoting the production of proinflammtory cytokine IL-6 and in decreasing the probability of inflammation-contributed cancer during the stage of acute inflammation.

Let-7 was one of the first miRNAs identified, and extensive research has focused on let-7 target prediction to uncover its biological role ^42^. Its family members are highly conserved across animal species with regard to seed sequences and have been reported to be important regulators of tumours and immunity, specifically in repressing the production of IL-6, a key proinflammatory cytokine during acute inflammatory response ^8^. We found that macrophages preferentially express let-7a to inhibit the production of IL-6 in acute inflammatory response. One key question regarding acute inflammatory response is how macrophages rapidly release the inhibitory effect of let-7 on IL-6 during pathogenic infection. Our analysis of the gene expression profiles of mouse peritoneal macrophages stimulated with LPS showed that *RasGRP1* mRNA may be the most important ceRNA for *IL-6* to sponge sufficient let-7a molecules to derepress the inhibitory effect of let-7 on the production of IL-6. Whether the other known targets of let-7, such as *Edn1*, *Socs1*, *Olr1* and *TNFaip3* ^43–47^, have synergistic effects with *RasGRP1* in regulating the expression of IL-6 during acute inflammatory response will require further investigation.

RasGRP1 is a Ras guanine nucleotide releasing factor that activates Ras and its downstream ERK and AKT signalling pathways ^48, 49^. However, the function of RasGRP1 in tumorigenesis has been debated ^50^. EGFR signalling-driven ERK and AKT signalling cascades are critical for the development and tumorigenesis of hepatocellular carcinoma, lung cancer, breast cancer and others ^51–53^. A recent study reported that RasGRP1 upregulation is associated with a better prognosis in colorectal cancer patients by limiting proliferative EGFR-SOS1-Ras-ERK signals ^40^. We found that overexpression of RasGRP1 restricts liver cancer cell growth by inhibiting proliferative EGFR-SOS1-Ras-AKT signals. However, whether RasGRP1-mediated EGFR-SOS1-Ras-ERK or -AKT signaling inhibition as a general mechanism also works in other inflammation-associated cancers may require further investigation.

In summary, our study demonstrated that RasGRP1 is essential regulator that promotes the inflammatory response by enhancing the production of IL-6, and restricts the growth of liver cancer cells by limiting the activation of EGFR-SOS1-Ras-AKT signalling. Under physiological conditions, especially during the stage of acute inflammatory response, RasGRP1 may play a key role in regulating inflammatory response and decreasing the probability of inflammation-contributed cancer.

### Materials and methods Mice, cells and reagents

Wild-type C57BL/6 mice (6–8 weeks of age) were purchased from Joint Ventures Sipper BK Experimental Animals (Shanghai, China). *IL6^−/−^* mice were purchased from the Jackson Laboratory. The mice were bred in pathogen-free conditions. All animal experiments were conducted in accordance with the principles of the Declaration of Helsinki and approved by the Medical Ethics Committee of Zhejiang University School of Medicine. HEK293 cells and RAW264.7 cells were obtained from the American Type Culture Collection (Manassas, VA) and cultured according to recommended protocols. Mouse peritoneal macrophages and human peripheral blood monocyte-derived macrophages (MDMs) were prepared and cultured as described previously ^30^. LPS (0111:B4) and poly (I:C) were purchased from Sigma (St. Louis, MO, USA). CpG ODN was obtained from Invivogen (San Diego, California). Antibody against RasGRP1 was from Abcam, Inc. (Cambridge, MA). Antibodies for JNK1/2, p-JNK1/2 (Thr183/Tyr185), ERK1/2, p-ERK1/2 (Thr202/Tyr204), p38, p-p38 (Thr180/Tyr182), AKT, p-AKT (Ser473), IkBα, p-IkBα (Ser32/36) and IKKα/β were purchased from Cell Signaling Technology (Beverly, MA). Antibody against β-actin was from Sigma-Aldrich.

### MiRNA mimics, sponges and inhibitors

Let-7a mimics (cholesterol modified oligonucleotides, labelled with FAM), let-7a sponges #1, #2, and #3 (cholesterol modified oligonucleotides, labelled with Cy3) and let-7a inhibitors (dsRNA oligonucleotides) from GenePharma (Shanghai, China) were used for the overexpression and inhibition of let-7a. Macrophages were transfected with miRNAs at a final concentration of 20 nM.

### Plasmid construction and transfection

The CDS and 3’UTR of *RasGRP1* and *IL-6* were obtained from mouse macrophage cDNA and cloned into the pcDNA3.1 vector. Each constructed plasmid was confirmed by sequencing. The corresponding primers are listed in the Supplementary information, Table S1. JetPEI transfection reagents (Polyplus Transfection, Illkirch, France) were used for cotransfection of plasmids and miRNAs into HEK293T cells.

### RNA interference

Small interfering RNA (siRNA) duplexes ^30^ were synthesized for the knockdown of mouse *RasGRP1* and human *RasGRP1* (Shanghai GenePharma Co., Shanghai, China). The siRNA duplexes of 5’-GCUCCAUCUAUUCCAAGCUTT-3’ (sense) and 5’-AGCUUGGAAUAGAUGGAGCTT-3’ (antisense) were used for mouse *RasGRP1* knockdown, whereas siRNA duplexes of 5’-GCGGGAUGAACUGUCACAATT-3’ (sense) and 5’-UUGUGACAGUUCAUCCCGCTT-3’ (antisense), were used for human *RasGRP1* knockdown. The siRNA duplexes of 5’-UUCUCCGAACGUGUCACGUTT-3’ (sense) and 5’-ACGUGACACGUUCGGAGAATT-3’ (antisense) were synthesized as RNA interference controls (Ctrls). The siRNA duplexes were transfected into macrophages using INTERFERin according to the protocol (Polyplus-Transfection, Illkirch, France).

### Quantitative PCR

Quantitative PCR analysis was performed with specific primer pairs (Supplementary Table S2) as previously described ^7^.

### Enzyme-linked immunosorbent assays (ELISAs) of cytokines

ELISA kits for human and mouse TNFα and IL-6 were obtained from R&D Systems. The concentrations of TNFα and IL-6 in the culture supernatants or serum were determined by ELISA as described previously ^30^.

### Western blotting

Total cell lysates were prepared as described previously ^30^. Cell extracts were subjected to SDS-PAGE, transferred onto nitrocellulose membranes, and subjected to immunoblot analysis as previously described ^30^.

### Gene microarray analysis

RNA was extracted from macrophages or LPS-stimulated macrophages using an RNA extraction kit (Fastagen Biotech Co., Shanghai, China). Gene expression microarray analysis was performed using the Mouse OneArray Plus® Microarray (Hsinchu, Taiwan) and performed according to the instruction manual. The changes in the let-7a target genes predicted by TargetScan (http://www.targetscan.org/vert_72) are presented in a heat map according to the microarray results.

### Liver cancer tissue microarray

A liver cancer tissue microarray contained 90 carcinoma tissues and 90 matched adjacent tissues was purchased from Shanghai Outdo Biotech Co. (Shanghai, China). Immunohistochemistry (IHC) analysis was performed using an anti-RasGRP1-specific antibody (Abcam, Inc. Cambridge, MA) according to the instruction manual. Briefly, the tissue sections were blocked with goat serum and then incubated with anti-RasGRP1 antibody overnight at 4 °C. The sections were stained with 3,3-diaminobenzidine and counterstained with haematoxylin after incubation with secondary antibody. PBS was used as a negative control. The staining intensity was used to examine the expression of RasGRP1 in liver cancer tissue.

### CRISPR-Cas9-mediated depletion of the 3’UTR of *RasGRP1*

The 3’UTR of *RasGRP1* was knocked out using the CRISPR-Cas9 system as described previously ^30^. The small guide RNA (sgRNA) sequences were designed by http://crispr.mit.edu as follows: sgRNA 1: 5′-TCTGAGGTAGCGAACACTGT-3′; sgRNA 2: 5 ′ -GGAAGGAGGCGGGCAAGTGA-3 ′ . These sequences were subcloned into the pLentiCRISPR V2 plasmid and cotransfected into RAW264.7 cells. After 2 µg/ml puromycin induction for 7 days, clones propagated from single cells were picked out. The depletion of the 3′UTR of *RasGRP1* was screened by PCR and confirmed by DNA sequencing.

### Dextran sulphate sodium (DSS)-induced colitis and histology

For establishment of the model of DSS-induced colitis in *IL-6*^−/−^ mice and in wild-type mice, all mice (8 weeks of age) received water containing 2.5% DSS for 7 days followed by normal water. On the indicated days, the body weight of each mouse was measured. On day 10 after DSS treatments, sera were collected for IL-6 ELISAs, the colon length was measured, and the colons were subjected to H&E analysis to determine the severity of inflammation ^30^.

### Statistical analysis

Data were analysed using GraphPad Prism 5. Results are given as the mean±s.e. or mean±s.d. Comparisons between two groups were performed using Student’s t-test or the Mann–Whitney U-test. Survival curves were generated using the Kaplan–Meier method and compared using a log-rank test. Box and whisker plots were compared by the Mann-Whitney test. Statistical significance was defined as a *p* value < 0.05.

### Data availability

The microarray dataset generating during the current study are deposited in NCBI’s Gene Expression Omnibus ^54^ and are accessible through GEO Series accession number GSE165800 (https://www.ncbi.nlm.nih.gov/geo/query/acc.cgi?acc=GSE165800)

## Acknowledgements

We thank Prof. Chen Liu (YaleUniversity) for providing the Huh7 cells. This study was supported by grants from the National Natural Science Foundation of China (81601383, 81971498, 81730064 and 31670877), the National Science and Technology Major Project (2017ZX10202201), the China Postdoctoral Science Foundation (2015M571888), the Fundamental Research Funds for the Central Universities and the Hunan Natural Science Foundation (2018JJ3091).

## Author contributions

S.T., J.W. and H.Z. designed the experiments and supervised the project; C.W., X.L., C.Y., L.W., R.D., H.L., Z.C., Y.Z. and S.F. performed the experimental work; S.T., H.S., J.W. and H.Z. analyzed results and wrote the manuscript.

---

**Fig. S1.**
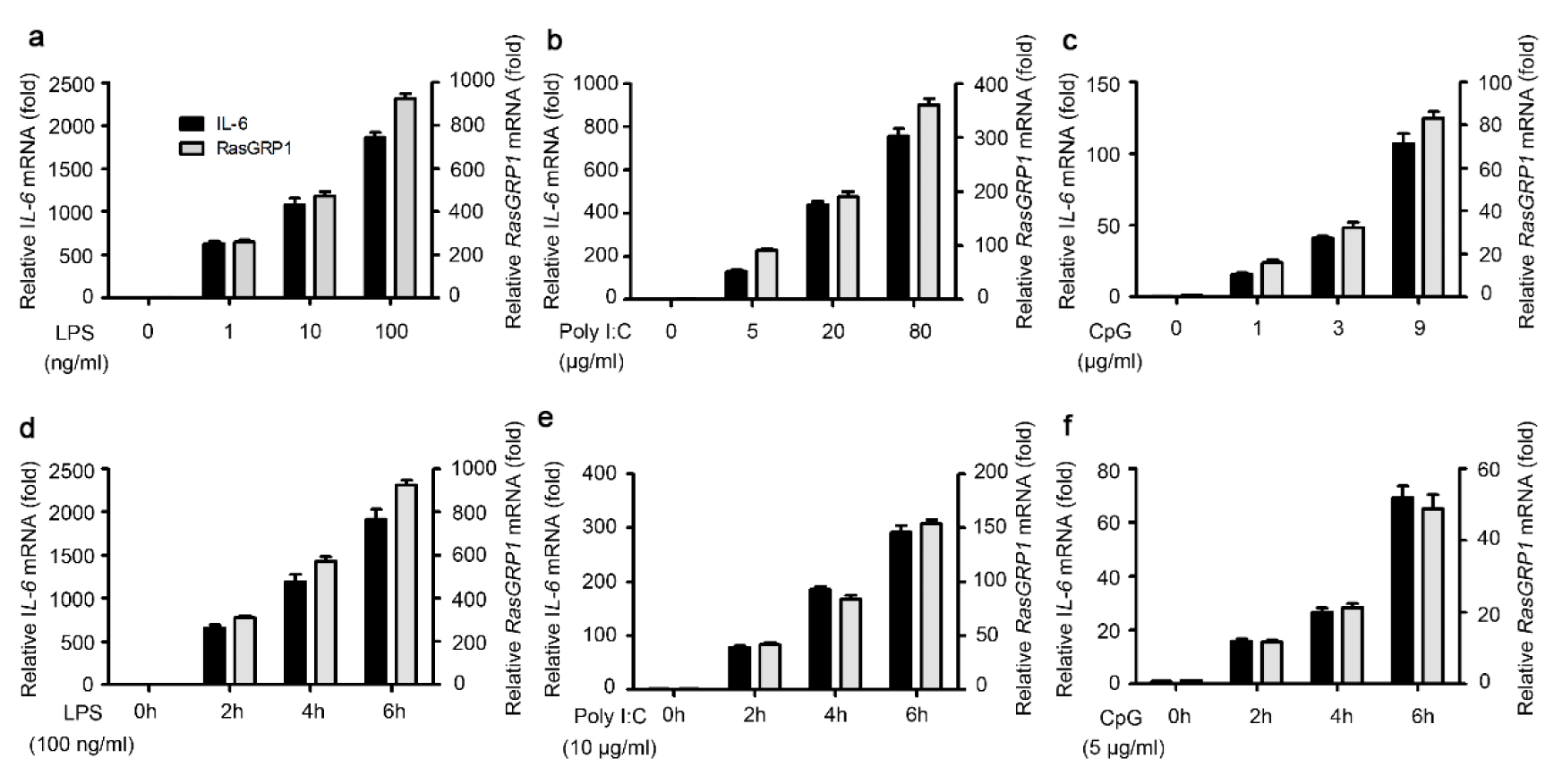
RasGRP1 co-expresses with IL-6 in acute inflammatory response. **a-c** Q-PCR analysis of IL-6 and RasGRP1 mRNA expression in bone marrow-derived macrophages treated with different doses of LPS (0, 1, 10 or 100 ng/ml) (**a**), Poly (I:C) (0, 5, 20 or 80 μg/ml) (**b**), or CpG ODN (0, 1, 3 or 9 μg/ml) (**c**) for 6h. **d-f** Q-PCR analysis of IL-6 and RasGRP1 mRNA expression in bone marrow-derived macrophages treated with 100 ng/ml LPS (**d**), 10 μg/ml Poly (I:C) (**e**), or 5 μg/ml CpG ODN (**f**) for the indicated hours. Data are shown as mean ± S.E.M. of three independent experiments (**a-f**).

**Fig. S2.**
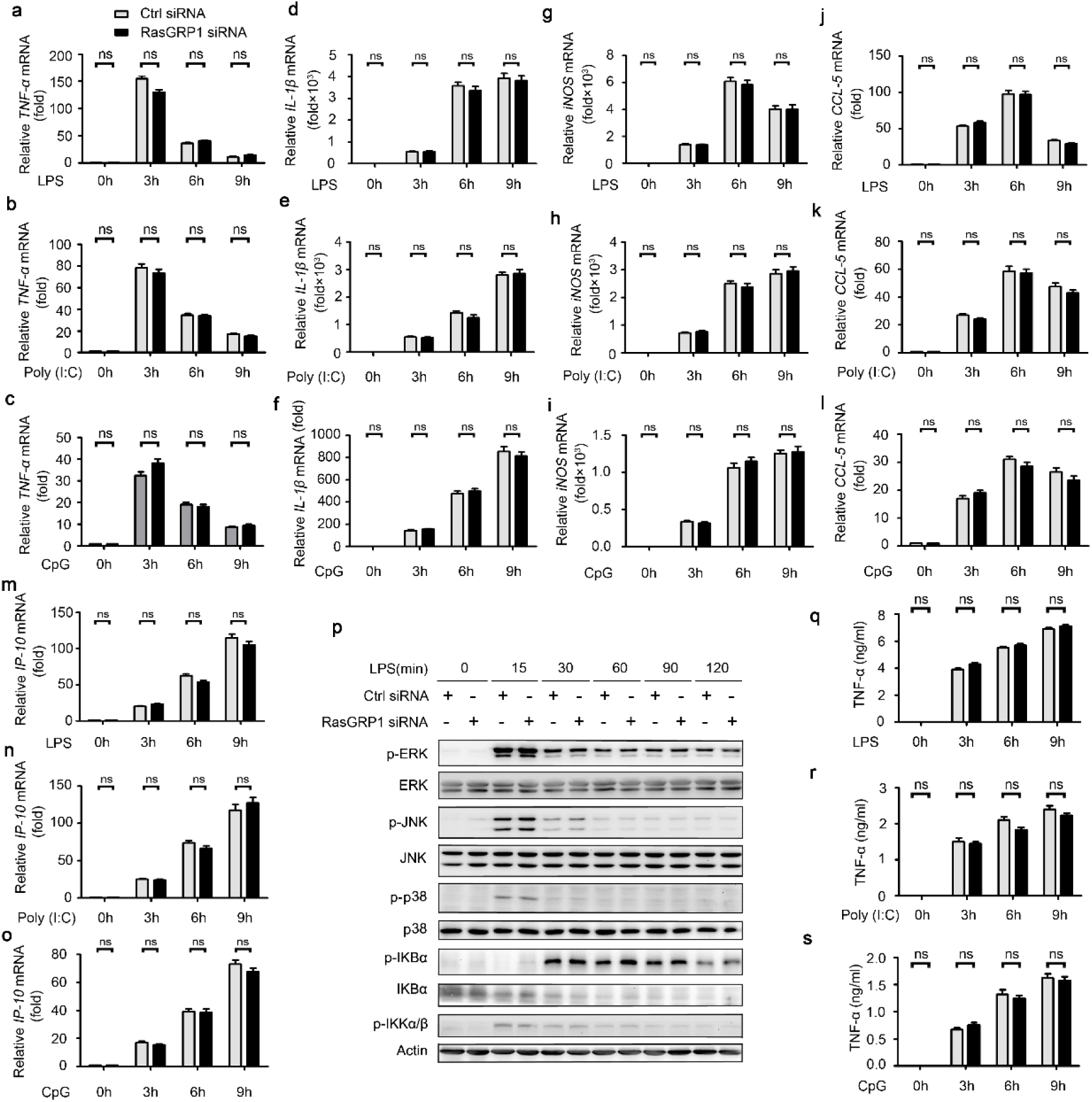
RasGRP1 silence has no effect on the mRNA level of cytokines in peritoneal macrophages. **a-o** Q-PCR analysis of TNF-α (**a-c**) IL-1β (**d-f**), iNOS (**g-i**) CCL-5 (**j-l**) or IP-10 (**m-o**) mRNA expression in peritoneal macrophages transfected with RasGRP1 siRNA and 48h later treated with LPS (**a, d, g, j, m**), Poly (I:C) (**b, e, h, k, n**) or CpG ODN (**c, f, I, l, o**) for 6h. **p** Immunoblot analysis of indicated molecules in lysates of peritoneal macrophages transfected with RasGRP1 siRNA and 48h later treated with 100 ng/ml LPS for the indicated minutes. **q-s** ELISA quantification of IL-6 in supernatants of macrophages treated as in (**a-c**). Data are representative of three independent experiments with similar results (means ± S.D. in **a-o** and **q-s**). ns, not significant.

**Fig. S3.**
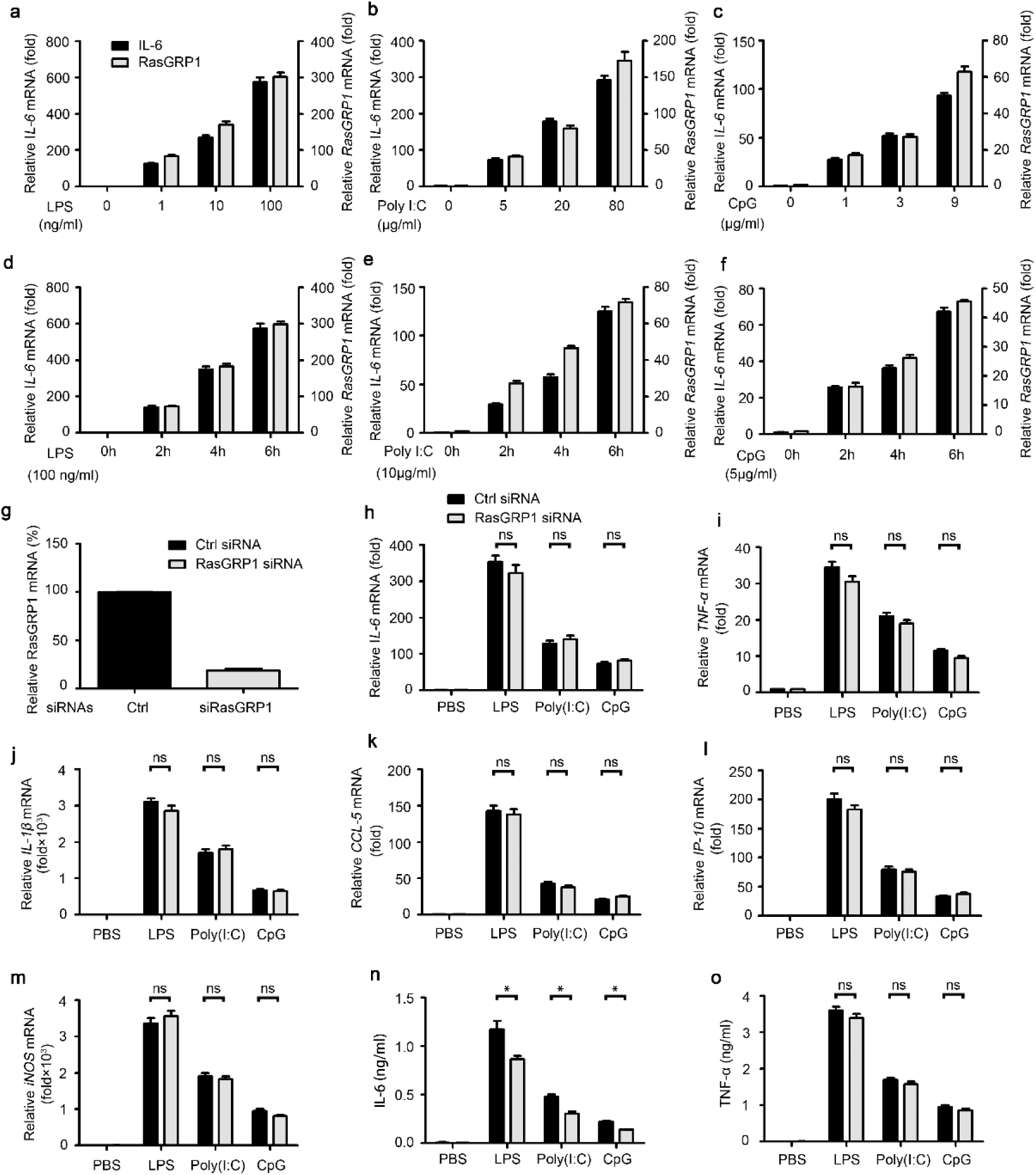
RasGRP1 co-expresses with IL-6 and RasGRP1 silence selectively inhibits IL-6 protein levels in human peripheral blood monocyte-derived macrophages. **a-c** Q-PCR analysis of IL-6 and RasGRP1 mRNA expression in human monocyte-derived macrophages treated with different doses of LPS (0, 1, 10 or 100 ng/ml) (**a**), Poly (I:C) (0, 5, 20 or 80 μg/ml) (**b**), or CpG ODN (0, 1, 3 or 9 μg/ml) (**c**) for 6h. **d-f** Q-PCR analysis of IL-6 and RasGRP1 mRNA expression in human monocyte-derived macrophages treated with 100 ng/ml LPS (**d**), 10 μg/ml Poly (I:C) (**e**), or 5 μg/ml CpG ODN (**f**) for the indicated hours. **g** Q-PCR analysis of RasGRP1 mRNA expression in human monocyte-derived macrophages 48h after transfection with RasGRP1 siRNA. **h-m** Q-PCR analysis of IL-6 (**h**), TNF-α (**i**) IL-1β (**j**), CCL-5 (**k**), IP-10 (**l**) or iNOS (**m**) mRNA expression in human monocyte-derived macrophages transfected with RasGRP1 siRNA and 48h later treated with LPS, Poly (I:C) or CpG ODN for 6h. ELISA quantification of IL-6 (**n**) and TNF-α (**o**) in supernatants of human monocyte-derived macrophages transfected with RasGRP1 siRNA and 48h later treated with LPS, Poly (I:C) or CpG ODN for 6h. Data are representative of three independent experiments with similar results. ns, not significant; * *p*<0.05.

**Fig. S4.**
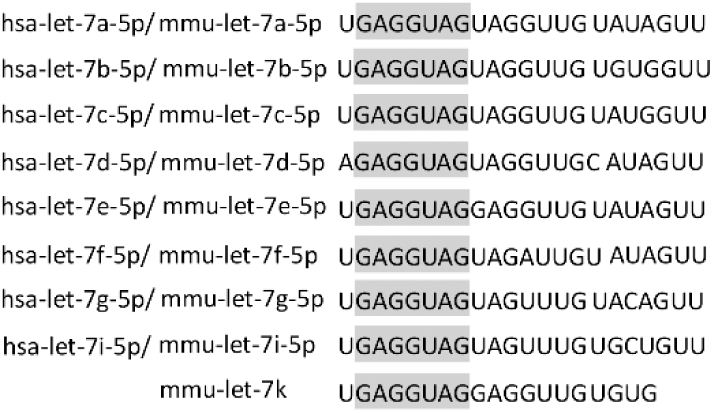
Let-7 family members have a conserved seed sequence. A schematic diagram showing let-7 family members with conserved seed sequences in humans and mice.

**Fig. S5.**
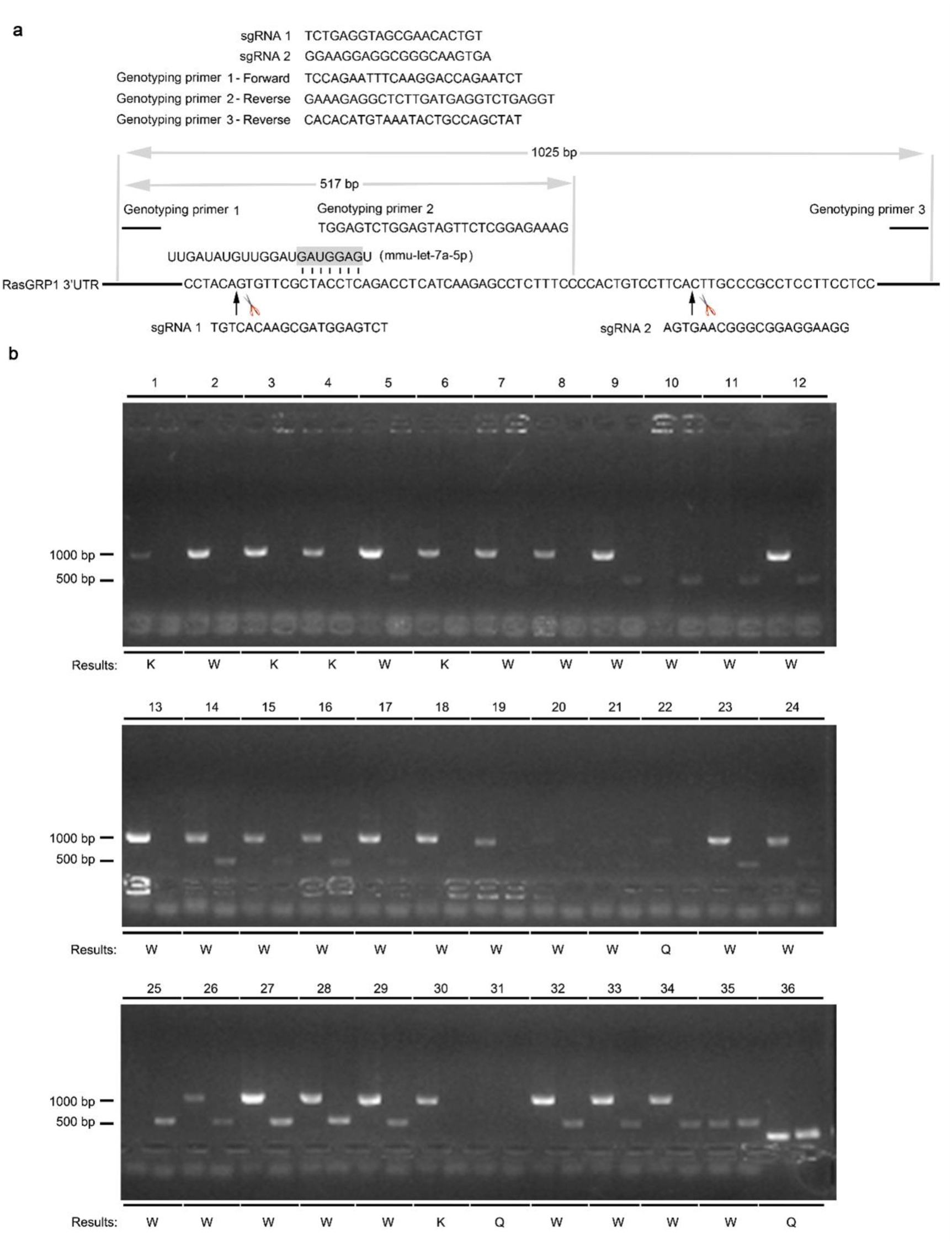
RasGRP1 3’UTR mutant clones constructed in RAW264.7 cells. **a** A schematic diagram showing the sequence of two sgRNAs designed for knocking out the binding site of let-7a on the RasGRP1 3’UTR and three genotyping primers designed for screening the targeted mutation. **b** PCR screening of the targeted mutation at the 3’UTR of the RasGRP1 gene in the RAW264.7 macrophage clone. The PCR results were evaluated as indicated at the bottom (“P” for positive, “W” for wild type, and “Q” for issues with DNA quality). Only the RAW264.7 macrophage clones showing positive results were selected and confirmed again by sequencing.

**Fig. S6.**
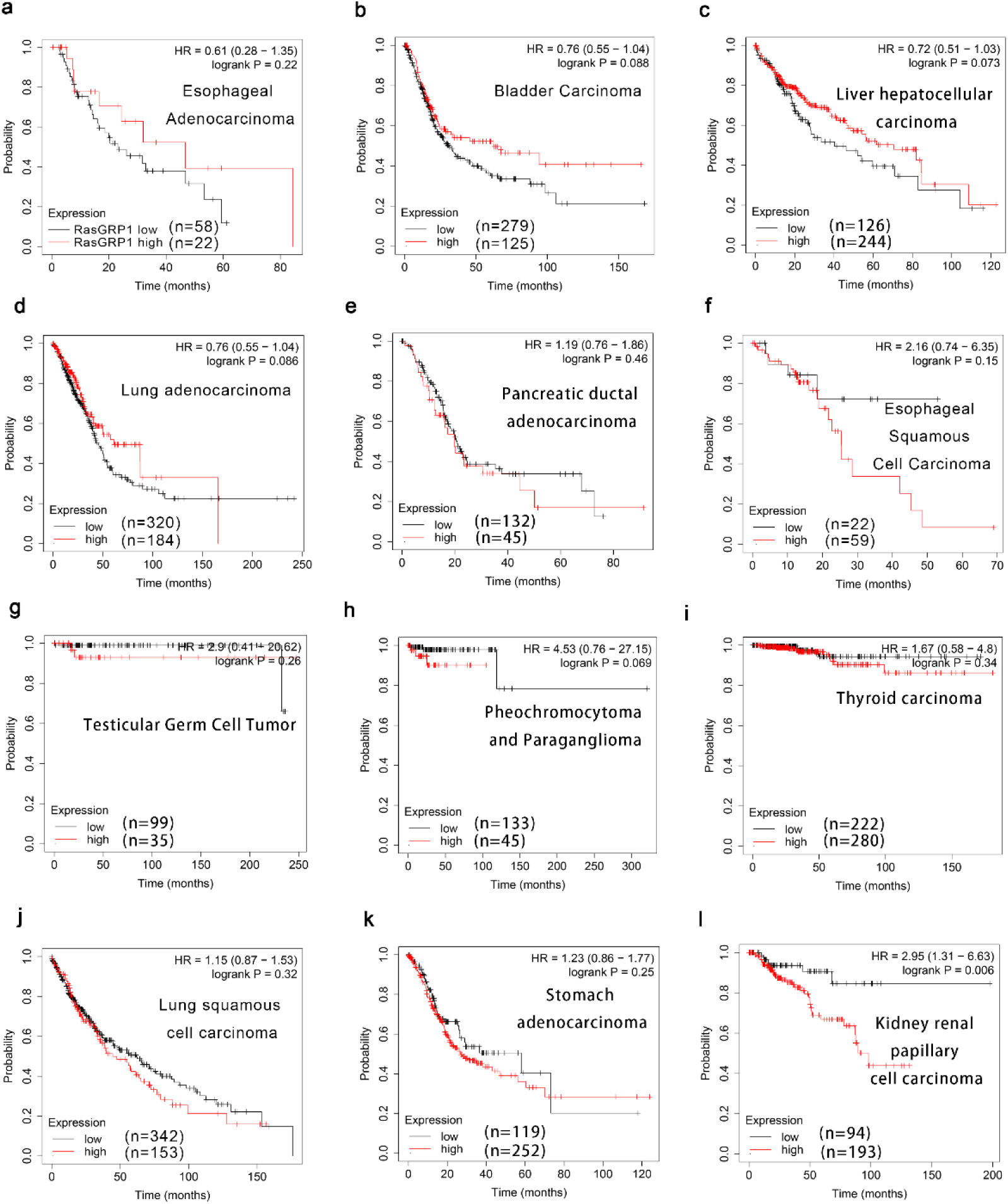
RasGRP1 expression correlates with the overall survival of cancer patients. **a-l** The survival curves from Kaplan-Meier plot profiles (http://kmplot.com/analysis/) for cancer patients stratified by high and low expression of RasGRP1.

**Supplementary Table S1.**
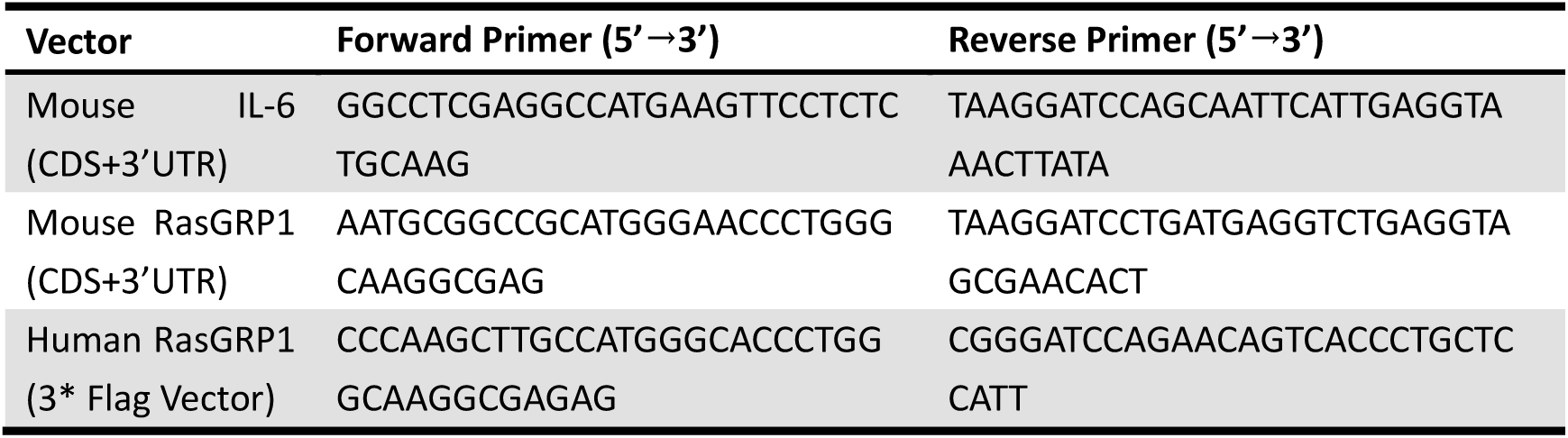
**Primers used in vector construction**

**Supplementary Table S2.**
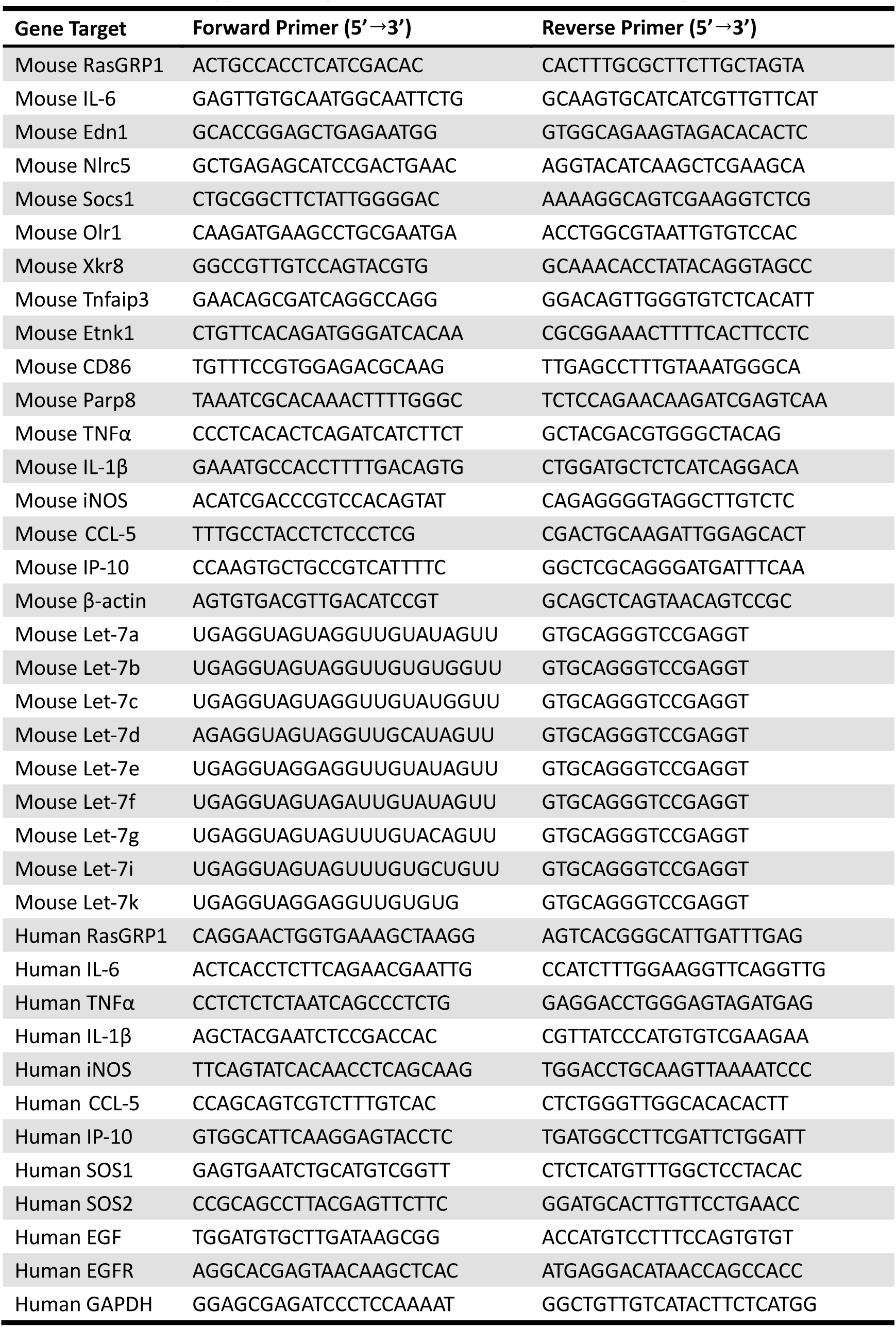
**Primers used in Q-PCR assays.**

